# Machine learning assisted mid-infrared spectrochemical fibrillar collagen imaging in clinical tissues

**DOI:** 10.1101/2024.05.22.595393

**Authors:** Wihan Adi, Bryan E. Rubio Perez, Yuming Liu, Sydney Runkle, Kevin W. Eliceiri, Filiz Yesilkoy

## Abstract

**Significance:** Label-free multimodal imaging methods that can provide complementary structural and chemical information from the same sample are critical for comprehensive tissue analyses. These methods are specifically needed to study the complex tumor-microenvironment where fibrillar collagen’s architectural changes are associated with cancer progression. To address this need, we present a multimodal computational imaging method where mid-infrared spectral imaging (MIRSI) is employed with second harmonic generation (SHG) microscopy to identify fibrillar collagen in biological tissues.

**Aim:** To demonstrate a multimodal approach where a morphology-specific contrast mechanism guides a mid-infrared spectral imaging method to detect fibrillar collagen based on its chemical signatures.

**Approach:** We trained a supervised machine learning (ML) model using SHG images as ground truth collagen labels to classify fibrillar collagen in biological tissues based on their mid-infrared hyperspectral images. Five human pancreatic tissue samples (sizes are in the order of millimeters) were imaged by both MIRSI and SHG microscopes. In total, 2.8 million MIRSI spectra were used to train a random forest (RF) model. The remaining 68 million spectra were used to validate the collagen images generated by the RF-MIRSI model in terms of collagen segmentation, orientation, and alignment.

**Results:** Compared to the SHG ground truth, the generated MIRSI collagen images achieved a high average boundary F-score (0.8 at 4 pixels threshold) in the collagen distribution, high correlation (Pearson’s R 0.82) in the collagen orientation, and similarly high correlation (Pearson’s R 0.66) in the collagen alignment.

**Conclusions:** We showed the potential of ML-aided label-free mid-infrared hyperspectral imaging for collagen fiber and tumor microenvironment analysis in tumor pathology samples.

## 1. Introduction

The tumor microenvironment (TME) is compositionally and structurally heterogeneous, hosting a complex network of biomolecules that encode a variety of biological signals, and metabolic and immune interactions. Collagen is the dominant structural protein in the extracellular matrix (ECM) of the TME^1^. Among the 28 types of collagen, fibrillar type I collagen forms a triple-helix structure and organizes itself into a fiber-like structure^2^, being the primary collagen in the ECM. It has been shown that the changes in the collagen (especially type I collagen) structure or distribution are linked to many diseases including cancer. Specifically, parameters like fiber density and orientation of collagen fibers have a significant impact on the progression and treatment of cancer^3–6^. For example, the stroma in the TME of pancreatic ductal adenocarcinoma (PDAC) is highly fibrotic, constituting up to 85% of the tumor volume^7^. PDAC fibrosis affects the efficacy of cytotoxic therapies and can compromise drug delivery^2^. Therefore, comprehensive analysis of fibrillar collagen in the TME is critical for understanding the role of collagen’s architectural changes in carcinogenesis and metastasis, which have implications for the development of effective cancer therapies and personalized treatments.

Conventional histological staining agents, such as Masson’s Trichrome, Movat’s Pentachrome, Van Gieson’s stain, and Picrosirius Red, can be used for collagen imaging ^2,8^. While these labeled techniques enable inspection of collagen in tissue samples using standard widefield microscopes, they are fairly laborious and time-consuming, requiring specialized protocols to overcome stain variation effects. Moreover, histological staining-based techniques are limited in resolving collagen fibers with high resolution and providing quantifiable metrics needed for prognosis and treatment studies. The implementation of artificial intelligence models to enhance the diagnostic potential of collagen-stained images has shown promise, but it is limited by the lack of insight into the biochemistry and the detailed morphology of the ECM^9,10^.

Label-free collagen-specific imaging modalities, such as polarized light^11,12^, and second harmonic generation (SHG) microscopy, have been demonstrated to overcome shortcomings of conventional stain-based imaging. Specifically, SHG is a second-order nonlinear coherent scattering process that is highly specific to non-centrosymmetric fibrillar collagen and its supramolecular fiber morphology. Besides, SHG microscopy has sub-micrometer resolution and can perform optical sectioning of tissue with an imaging depth of up to hundreds of micrometers^13,14^. SHG has been used to study tumor-associated collagen signatures^3,15^ (TACS). In particular, TACS-3, a pattern exhibiting high fiber alignment perpendicular to tumor boundaries, has been shown to be a negative prognosis factor in breast cancer^16^. Moreover, similar morphological signatures have also been found in other types of cancer such as skin^17,18^, ovarian^4,19^, prostate^20^, and pancreas^6,21^. While SHG imaging has elucidated the biomedical consequences of architectural changes in TME, the molecular mechanisms that drive collagen alterations are still poorly understood. Thus, there is a need for a multimodal approach where SHG and chemical imaging methods sensitive to molecular changes are employed together in TME investigations.

Infrared absorption spectroscopy is a label-free analytical technique that provides quantitative biochemical information via probing the vibrational bands of functional biomolecules. Specifically, in the mid-infrared (MIR) fingerprint region (∼ 800 cm^-1^ - 1800 cm^-1^), spectrochemical analysis of measured transmittance through a specimen reveals compositional information. Recently, tunable quantum cascade lasers (QCL) with high spectral power output enabled the development of wide-field MIR spectral imaging systems that can operate at room temperature with compact footprints^22–24^. The QCL-based MIR spectral imaging (MIRSI) can rapidly collect hyperspectral datasets from whole-slide tissue sections and has the unique capability to combine spatial and chemical information^25^. Previous important studies revealed that the MIR spectral analysis can detect different collagen types^26,27^, identify fibrosis in numerous tissues, including liver^28^, heart^29^, and bone marrow^30^, and provide critical prognostic information based on reactive stroma in TME-based investigations^31,32^. In these past reports, the MIRSI-identified fibrotic regions were primarily referenced to stained images of adjacent tissue sections. However, an objective benchmarking of MIRSI-detected collagen fibers with respect to the ground truth SHG microscopy on the same tissue sample has not been performed.

Here, we present a new label-free multimodal imaging approach using MIRSI and SHG imaging modalities to sequentially acquire complementary chemical and morphological information from the same biological tissue sample (Fig. 1). We first developed a protocol that enabled reliable image acquisition from the same tissue section using two different microscopes, which employ two distinct frequency regions of the electromagnetic spectrum, i.e., visible-near-infrared (λ= 890 nm) and mid-infrared (λ=5-10 µm). To classify fibrotic regions in pancreatic tissue samples based on collagen’s spectral signatures, we trained a Random Forest (RF) model using large hyperspectral MIRSI datasets and SHG images of the same tissues as the ground truth. This RF model, which we named RF-MIRSI, was then used to identify regions of high collagen composition from the whole-tissue sections. Finally, we both validated the RF-MIRSI fibrotic collagen segmentation method, referencing our findings to the SHG images. The RF-MIRSI method achieved a high average boundary F-score (0.8 at 4 pixels threshold) in the collagen distribution, high correlation (Pearson’s R 0.82) in the collagen orientation, and similarly high correlation (Pearson’s R 0.66) in the collagen alignment. Our label-free multimodal collagen fiber imaging approach is a key step towards future more comprehensive tumor tissue investigations where both morphometric and chemometric information are considered to better study tumor-promoting ECM alterations.

**Figure 1.**
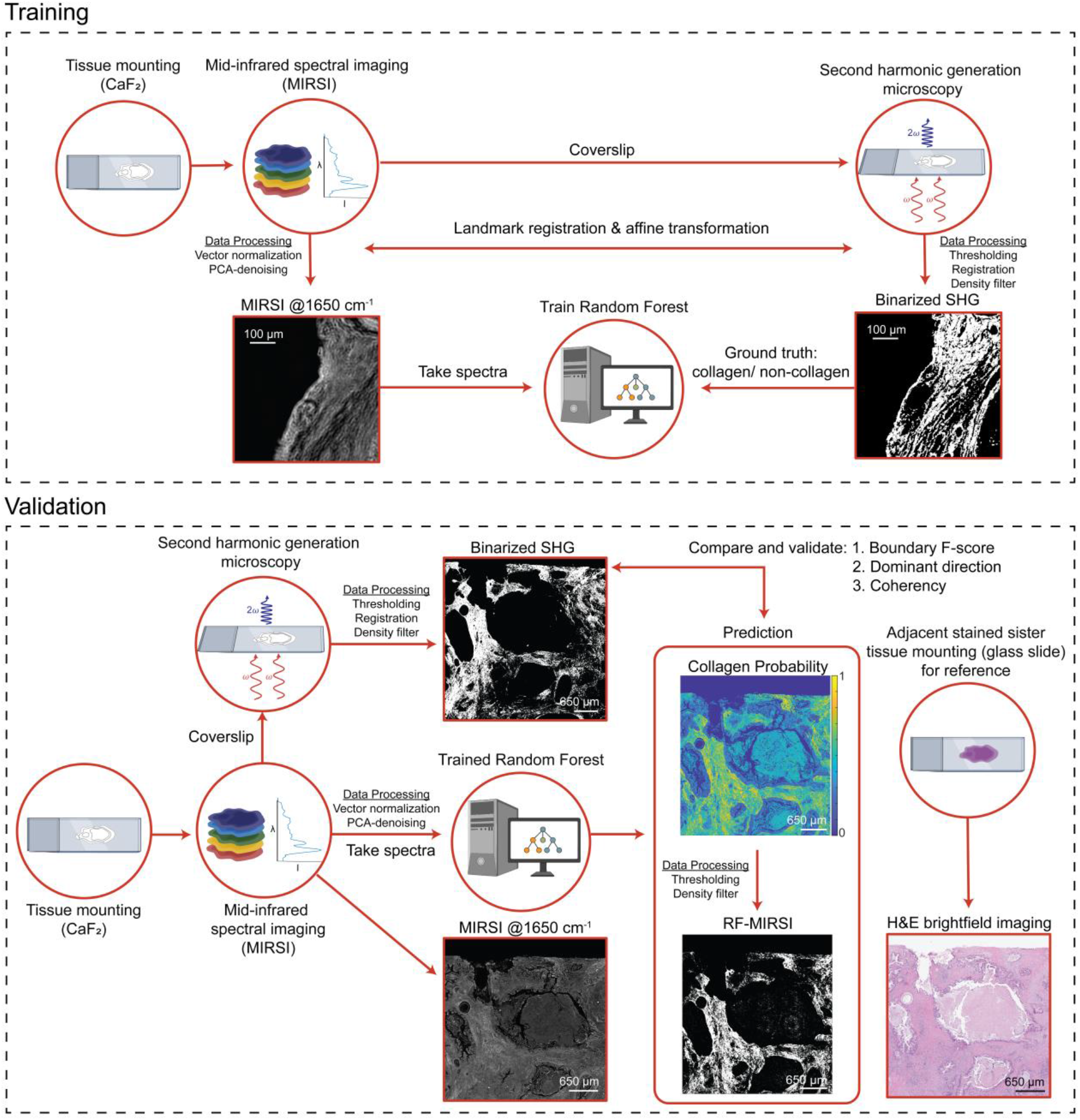
Schematic diagram of the workflow for label-free multimodal imaging using MIRSI and SHG: Random forest training workflow. **(top)**: A pancreatic tissue section is first mounted onto an infrared-transparent CaF_2_ substrate for MIRSI imaging. Subsequently, the tissue section is enclosed with a coverslip and imaged with a SHG microscope. SHG images are used as ground truth for the RF model training after the registration of MIRSI and SHG images. **Validation workflow (bottom):** Once the RF model is trained, an unused independent subset of the image data is used to validate the RF-MIRSI results. To generate SHG-like images from the RF-MIRSI predicted collagen pixels, further image processing, such as thresholding and density filtering, is conducted. Finally, RF-MIRSI images are compared to SHG images that have also undergone further processing, such as thresholding and density filtering, for equivalent comparison. Adjacent tissue slices are stained with H&E and imaged on a standard brightfield microscope for reference. Scale bars are 100 µm for both images in the training section and 650 µm for all images in the validation section. SHG: second harmonic generation; MIRSI: Mid-infrared spectral imaging; RF: Random Forest

## 3. Results

### 3.1 RF training and RF-MIRSI

MIRSI data of 5 tissues were collected, constituting ∼350 ROIs of 480x480 pixels or around 80 million spectra. To illustrate our raw MIRSI spectral data, Fig. 2 shows a typical unprocessed MIRSI image collected at 1650 cm^-1^ illumination that corresponds to the protein amide I band along with 10 randomly selected spectra for each of the 3 outlined ROIs. The example MIRSI image consists of four (2x2) stitched ROIs collected from a pancreatic tissue section. To train the RF, 1.4 million spectra, including collagen and non-collagen pixels, were used. These training spectra were pre-processed to exclude non-tissue regions and reduce noise. For the ML training, SHG image pixels were used as classification labels. MIRSI images are registered to SHG images using landmark registration, which matches their scaling, translation and rotation. Subsequently, the trained RF model was applied to the rest of the spectra collected from the tissues. For each pixel-associated spectrum, the RF outputs the probability of how likely that pixel contains collagen. More details about the data pre-processing workflow and the training of the RF can be found in the Materials and Methods.

**Figure 2.**
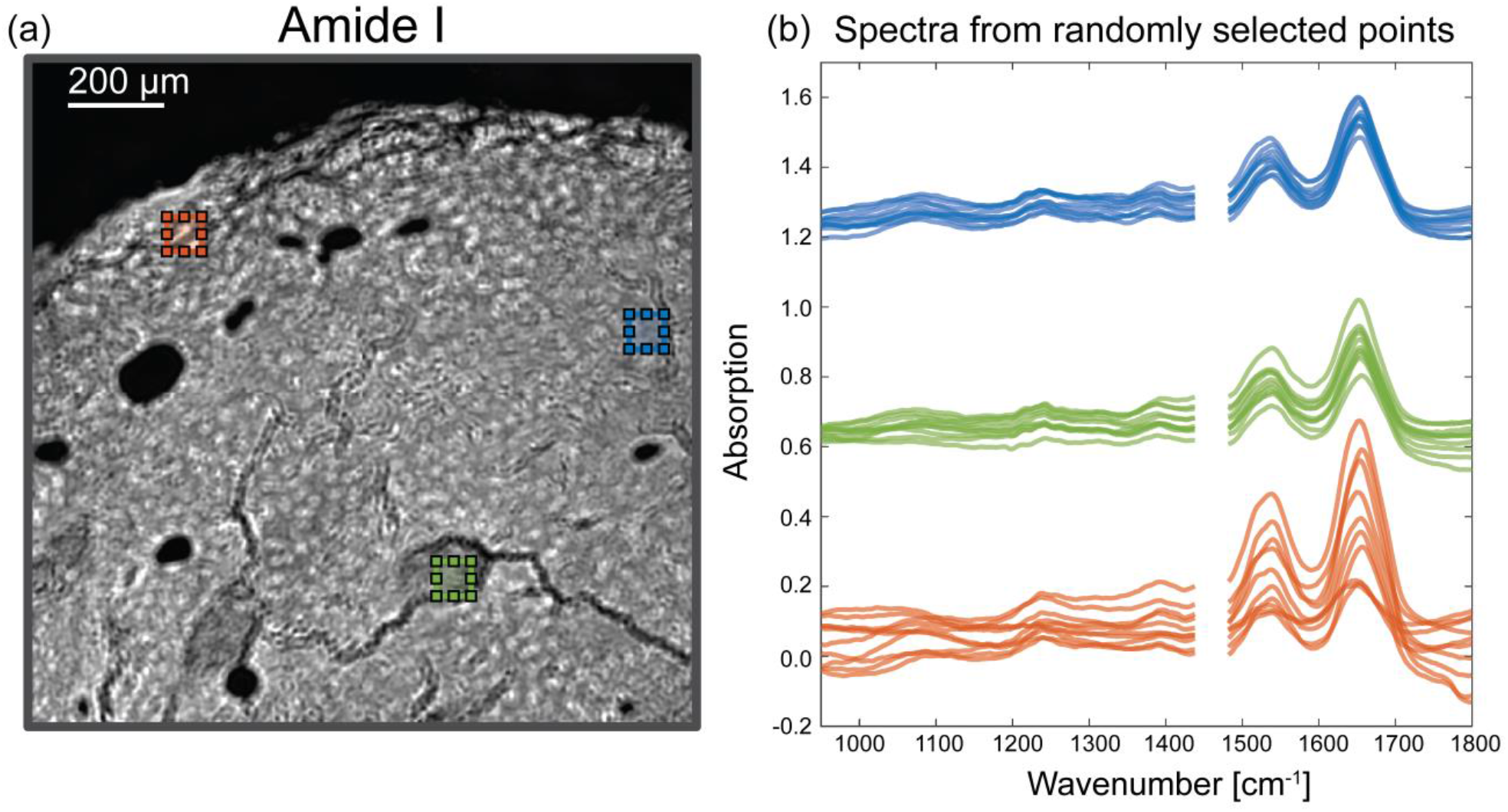
A representative MIRSI data. (a) MIR image collected at the 1650 cm^-1^ protein amide I band, (b) The MIR spectra of each of ten randomly chosen pixels from three ROIs in a pancreatic tissue section. The data from the spectral range of 1460 cm^-1^-1480 cm^-1^ was omitted due to the QCL switching issues with our instrument.

### 3.2 Correlation between binarized RF-MIRSI and SHG images

Fig. 3 presents the results of our correlation investigation between the RF-MIRSI-predicted collagen pixels and the SHG ground truth. A representative MIRSI image of a tissue region collected at 1650 cm^-1^ protein amide I band illumination is shown in Fig. 3a. The RF-MIRSI predicted collagen pixels from the same tissue region are shown in green in Fig. 3b. To qualitatively illustrate correlation between RF-MIRSI-predicted and SHG-identified collagen pixels, Fig. 3b shows the binarized SHG image in pink and the pixels that are identified as fibrillar collagen by both methods in white. Collagen fiber’s dominant direction and coherency were also calculated in a subregion, outlined with a white box in Fig 3b, and the findings were presented in Fig. 3c. For reference, an unprocessed SHG image (raw) counterpart of the same subregion is shown in Fig. 3d.

**Figure 3.**
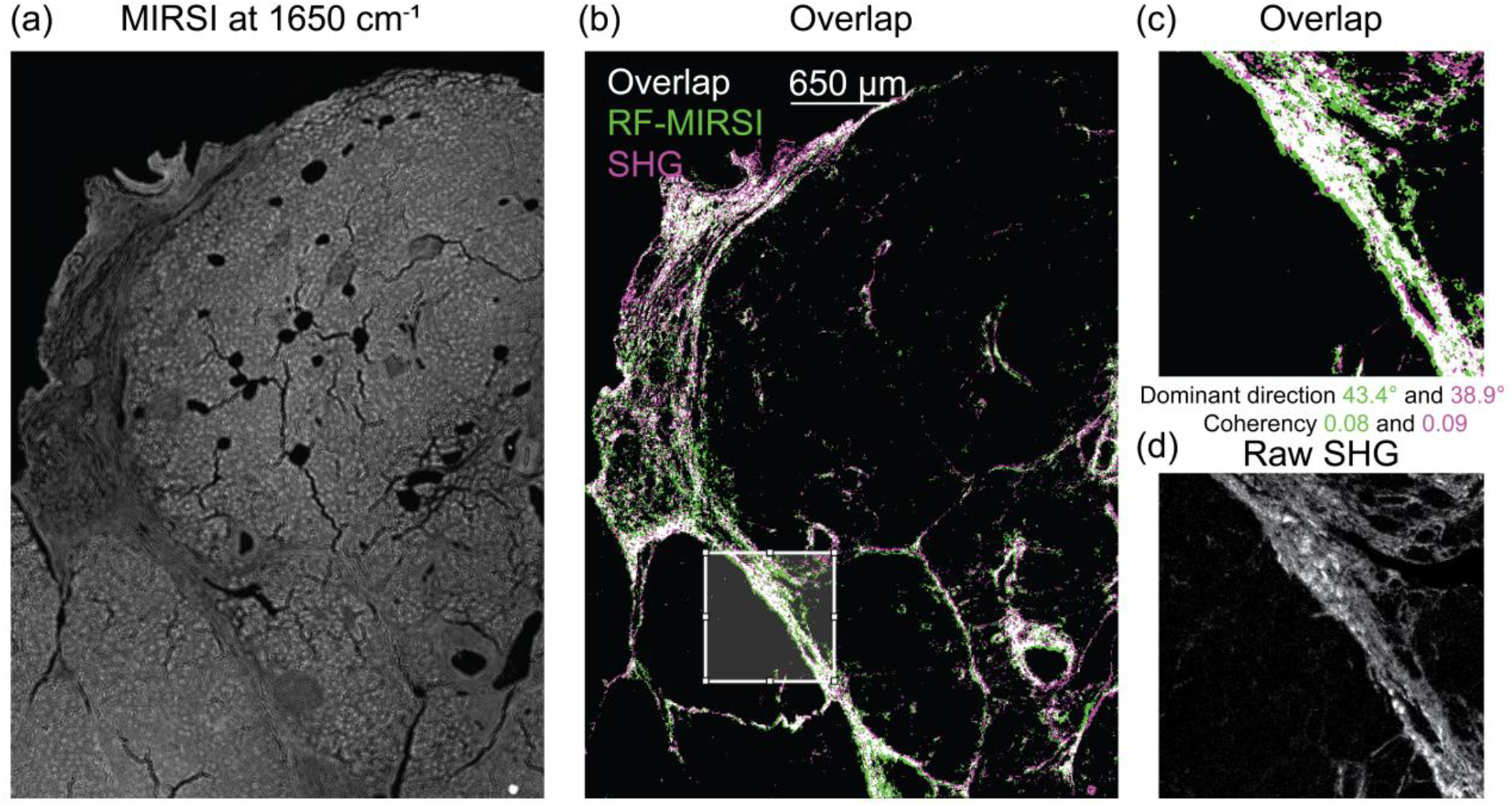
Correlation between RF-MIRSI and SHG images. (a) MIRSI image collected at 1650 cm^-1^ protein amide I band of a representative tissue, (b) overlay of corresponding RF-MIRSI-predicted collagen image (green) and SHG image (pink), where overlapping pixels (white) indicate correspondence in between. (c) shows a subregion enclosed with the white box from (b) presenting calculated dominant direction and coherency. (d) raw SHG image of the same region in (c) is shown for reference. The scale bar of length 650 μm shown in (b) is valid for (a) and (b). (c) and (d) are both 650 x 650 μm^2^ in size.

For the quantitative evaluation of our multimodal imaging technique, we used BF-score, dominant angle, and coherency as metrics. We first divided RF-MIRSI data from 5 tissues into 33 large ROIs of various sizes (all ∼mm^2^). The average BF-scores calculated for each of the 33 ROIs are shown in Fig. 4a for various pixels and collagen probability thresholds. For the dominant direction and coherency calculations, we used smaller ROIs (480x480 pixels) to preserve granularity. The Bland-Altman plot shown in Fig. 4b depicts the agreement of the dominant direction for both SHG and RF-MIRSI measurements with Pearson’s R of 0.82. Fig. 4c depicts the distribution of the absolute angle difference between the two techniques with a mean of -5.8 degrees and a standard deviation of 14.9 degrees. The coherency calculated from both techniques are shown in Fig. 4d with Pearson’s R of 0.66.

**Figure 4.**
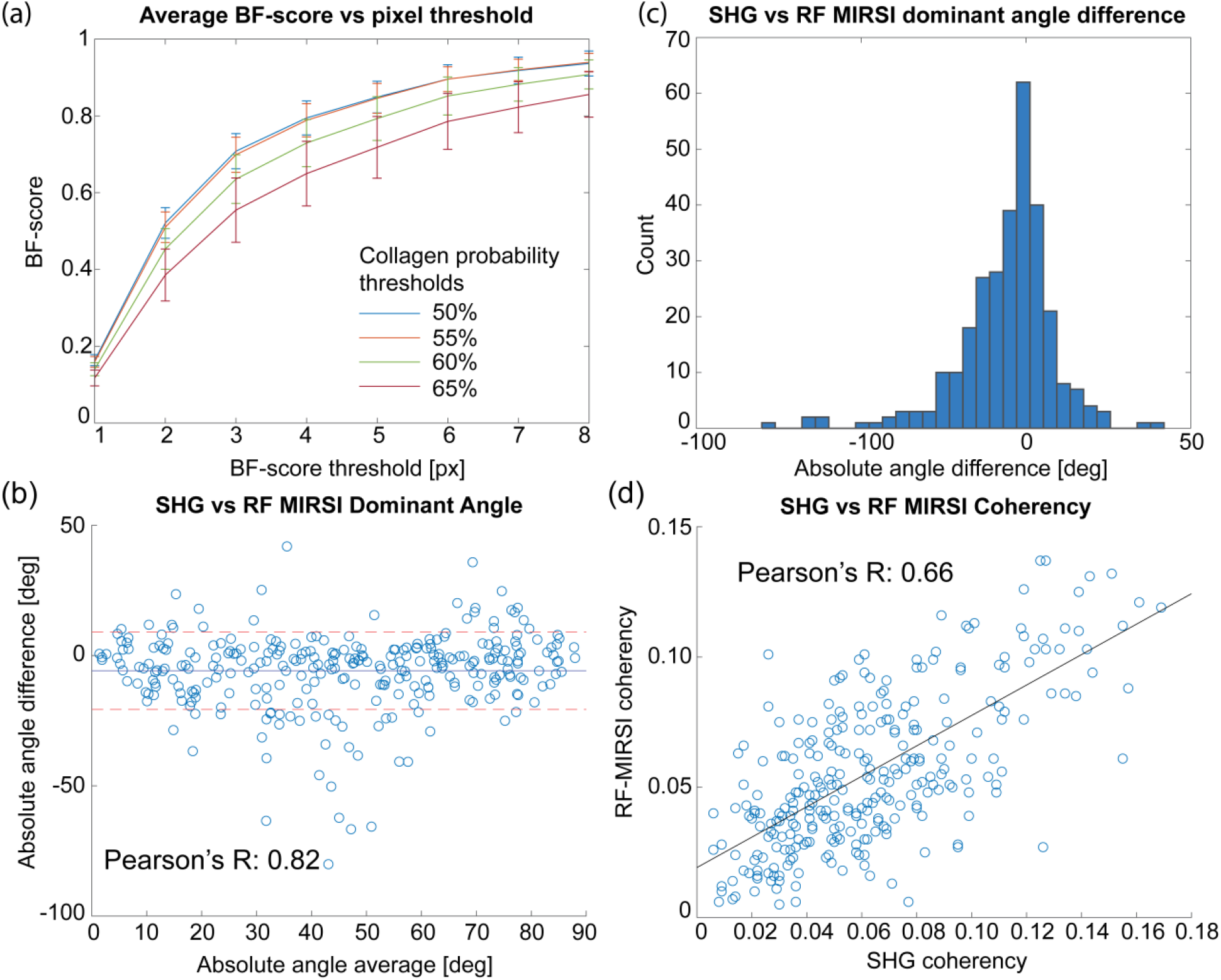
Quantitative validation of RF-MIRSI-identified collagen based on ground truth SHG images. (a) BF-score calculated to validate RF-MIRSI collagen prediction accuracy based on binarized SHG using various pixel and collagen probability thresholds. Error bar indicates standard deviation. (b) Bland-Altman plot comparing the dominant direction of collagen calculated using OrientationJ for both RF-MIRSI collagen probability and SHG images along with its Pearson’s R value. (c) The distribution of the calculated absolute angle difference. (d) Alignment (coherency) calculation between SHG and RF-MIRSI along with its Pearson’s R-value.

### 3.3 Wavenumbers significant in detecting collagen as identified by Random Forest

Fig. 5a shows the wavenumbers as a function of their importance in detecting collagen as identified by the RF model. The top 20 best predictor wavenumbers are colored in maroon. To further examine this, the average and standard deviation of collagen and non-collagen spectra from the training data are shown in Fig. 5b with the top 20 wavenumber predictors indicated with stars.

**Figure 5.**
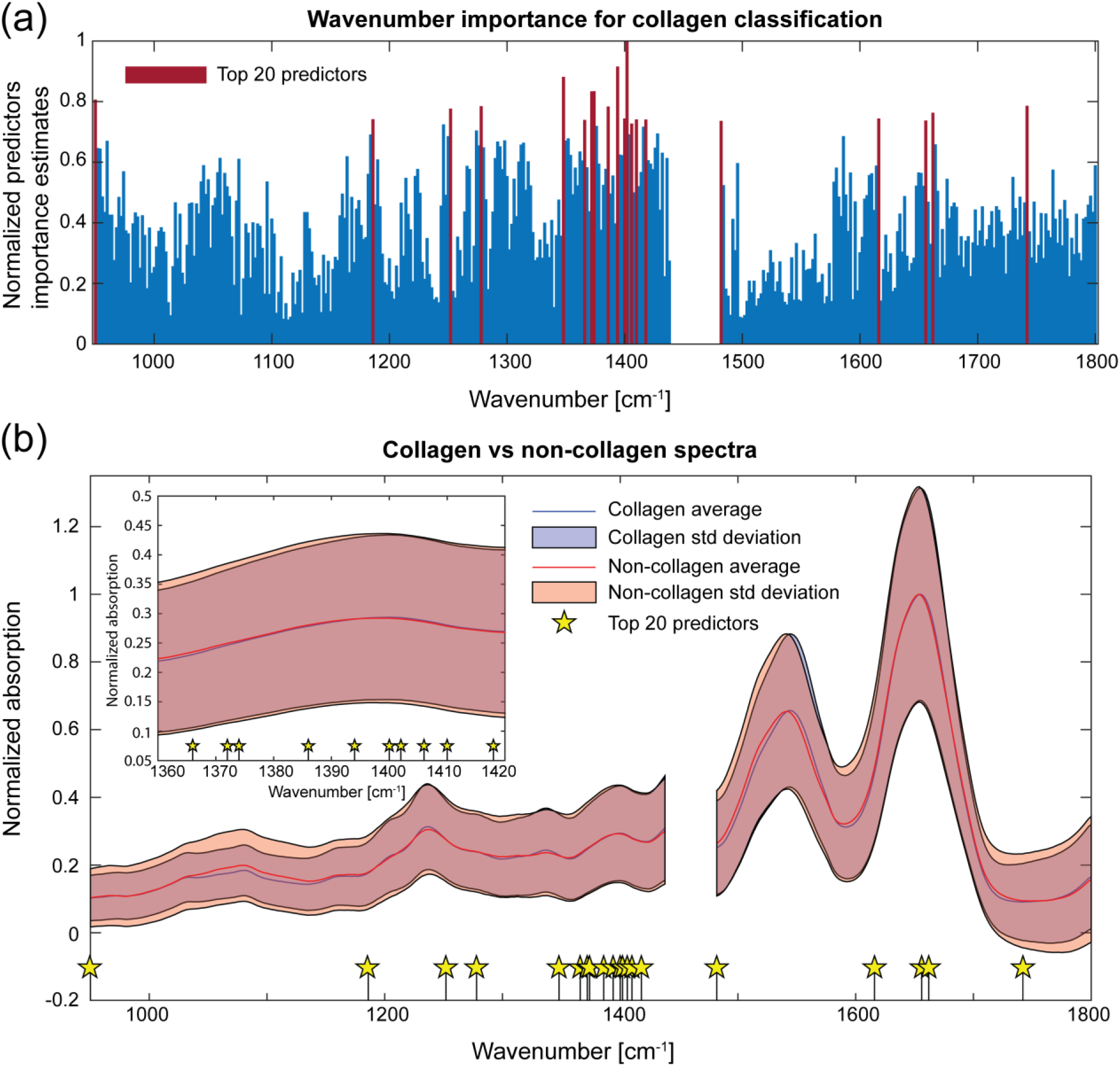
Importance of each spectral wavenumber identified by the RF model for collagen prediction. (a) The predictor importance values at each wavelength; the top 20 most important predictors are highlighted in maroon. (b) Average collagen and non-collagen spectra from the training dataset, together with the average of its standard deviation (shaded area).

## Discussion

In cancer, the growth of fibrous tissue around tumor (also referred to as a desmoplastic reaction) has been shown to be an important hallmark of TME, where it presents distinct similarities to the wound healing response^45–47^. Previously, highly organized fibrillar collagen patterns in TME were identified as negative cancer prognostic markers using SHG imaging, which is the current “gold-standard” in fibrillar collagen studies^6^. Despite their important prognostic potential, stromal-based biomarkers are not yet part of the current clinical histopathology because SHG is a low-throughput imaging technique and requires extensive expertise and costly instrumentation. MIRSI is a label-free and rapid hyperspectral digital tissue imaging technique, which can collect whole-slide tissue images within minutes. Here, we trained a machine learning algorithm using the gold-standard SHG images to generate fibrillar collagen maps using the rapid and high-throughput MIRSI modality. Our qualitative (Fig. 3) and quantitative (Fig. 4) investigations revealed that RF-MIRSI-predicted fibrotic collagen tissue regions correlate well with the gold-standard SHG imaging. Therefore, our method has the potential to contribute to the histopathology workflow by providing fibrillar collagen maps of whole-tissue pathology sections.

Notably, we report a significant difference between a MIR image of a tissue section collected at a single wavelength illumination associated with the protein band (Fig. 3a) and the collagen map predicted by the RF-MIRSI (Fig. 3b). This highlights the strength of our RF-MIRSI method in parsing out most of the tissue regions as non-collagen areas, even though they contain significant amounts of protein as evidenced in Fig. 3a, the amide I absorption signal. Moreover, contrary to traditional single-channel spectrometry, hyperspectral imaging modalities can collect spatially and spectrally rich datasets. When evaluated using machine learning and artificial intelligence models, MIR hyperspectral imaging can provide versatile information that is critical in the biomedical field.

In multimodal imaging, accurate image registration is inherently challenging, especially when images are collected by different instruments with distinct spatial resolution limits, as in the case of our SHG/MIRSI approach. To tackle the unavoidable image registration errors, we used the BF-score instead of the standard F-score as a validation metric. The average BF-score is calculated for all 33 ROIs encompassing all 5 tissues with various pixel and collagen probability thresholds as shown in Fig. 4a. Our results show that the average BF-score reaches ∼0.8 within 4 pixels threshold for collagen probability threshold of 50% and 55%, which indicates a good correlation between both sets of images. Below 4-pixels threshold, the average F-score is lower due to the misalignment in the registration of the images. The misalignment sources can include: i) human error while picking common registration landmarks for registration, which is amplified because SHG and MIRSI images are collected at different magnifications (20X vs 12X); and ii) mismatching non-affine deformations that come from the different imaging system such as barrel distortion as well as the cover slipping process for the SHG imaging (sample for MIRSI imaging is uncovered). While different factors impact the multimodal image alignment accuracy, using 4-pixels threshold (equivalent to ∼5 μm, which is close to the spatial resolution limit of MIRSI) in BF-score calculation successfully addresses this issue. Apart from misalignment, other factors that contribute to the deviations between the SHG and MIRSI results can also be explained by either the inherent differences between the two processes (SHG vs. absorption), such as their cross-sections and depth-of-field, or the difference in our implementation (SHG with circularly-polarized and MIRSI with linearly-polarized light).

Moreover, our quantitative validation results in Fig. 4a show that the average BF-score decreases with increasing collagen probability threshold. This can be explained by the lower true positive classification outcomes when a larger threshold is applied to RF-MIRSI images. Therefore, in this work, we first detected as many collagen-classified pixels as possible using a 50% collagen probability threshold, then minimized the non-structural collagen using a density filter (see Materials and Methods). A similar strategy and threshold were used while binarizing the SHG images.

We also compared the morphology in the RF-MIRSI collagen maps and SHG images via the dominant angular direction parameter calculated in individual ROIs (Fig. 4b) and identified a high correlation (Pearson’s R of 0.82). This is also evident in Fig. 4c, where the histogram distribution of the absolute angle differences between the imaging modalities peaks close to zero. Furthermore, the alignment (coherency) calculated for both techniques showed a high degree of correlation (Pearson’s R 0.66, Fig. 4d). Together, these results indicate that the RF-MIRSI images generated from MIRSI spectral data correlate well with the SHG images.

Among the ML models, RF has the advantage of quantifying the importance of data features in the decision-making process. We calculated the wavenumber importance by quantifying the increase in error generated by excluding a specific spectral feature (see Materials and Methods). Fig. 5a shows the 20 highest-ranked predictors. In MIRSI-based collagen studies, the spectral focus is usually on the protein-associated amide I and amide II bands^48–55^. Our RF model also identified four features falling within the amide I 1600-1700 cm^-1^ range among the top 20 highest ranked. Interestingly, the RF model heavily relied on the spectral region between 1360 cm^-1^ to 1420 cm^-1^, with 10 out of the top 20 predictors residing there, even though the average collagen and non-collagen spectra are not different in that region (Fig. 5b). A small window (1360–1340 cm^-1^) within this region contains the wagging vibration of the proline side chains present in type I collagen, found in biological tissues^52^. Therefore, the dominant dependence of the RF model on this spectral region must be due to the abundance of proline and 4-hydroxyproline in collagen triple-helix (∼22% occurrence of each in type I collagen^56^). This underscores the critical role of our holistic TME analysis via multimodal imaging, as it can provide access to biochemical information from the structurally altered tissue regions.

In future studies, specific biochemical information such as the integrity of collagen’s triple helix structure^48^, cross-linking collagen concentrations^48,53^, the collagen quality associated with non-enzymatic cross-linking^55^, and many more leveraging existing databases and literature^57^ can be investigated in the context of diseases. Such investigations can help elucidate the molecular drivers behind the morphological alterations in the TME observed in various cancer grades. Similarly, MIRSI can be used to complement SHG by analyzing interactions of collagen with other important ECM molecules such as fibronectin, which cannot be detected by SHG^58^.

Our results can be further improved by employing recently developed advanced laser scanning based MIRSI methods or photothermal imaging^59,60^, which can achieve a higher spatial resolution and better match SHG imaging^61^. Moreover, metasurface-enhanced MIRSI can be used to improve the sensitivity and selectivity of the absorption spectra by benefiting from the light-matter interactions at the photonic cavities with resonances tuned to the region with a high density of important predictors^62^.

In conclusion, the proposed RF-MIRSI model was successfully used to detect fibrillar collagen based on the MIRSI spectral data from pancreatic tissue samples. This technique can be adapted to other tissue types and can complement state-of-the-art imaging modalities and analytical techniques to further investigate the complex nature of fibrillar collagen in the TME.

## Acknowledgement

F.Y. acknowledges financial support from the National Institutes of Health (NIH) (grant #s: R61CA281795 and R21EB034411). KWE also acknowledges financial support from NIH (grant #s: R01CA238191, U54CA268069 and P41GM135019). B.R.P. acknowledges financial support from the UW-Madison College of Engineering Graduate Engineering Research Scholars (GERS) Fellowship. Moreover, the author(s) thank the University of Wisconsin Translational Research Initiatives in Pathology laboratory (TRIP), supported by the UW Department of Pathology and Laboratory Medicine, UWCCC (P30 CA014520) and the Office of The Director-NIH (S10 OD023526) for use of its facilities and services.

## Notes

### Competing Interest Statement

The authors have declared no competing interest.

